# LytF contributes to pilus extrusion during natural competence in *Streptococcus sanguinis* SK36

**DOI:** 10.1101/2025.09.04.674207

**Authors:** Rebekka Moe, Katarzyna Wiaroslawa Piechowiak, Leiv Sigve Håvarstein, Morten Kjos, Daniel Straume

## Abstract

Streptococci may enter a physiological state called competence, during which they express a specific set of genes required for exogenous DNA uptake and its subsequent integration into the genome through homologous recombination. This process, termed natural transformation, facilitates the horizontal acquisition of genetic material, potentially conferring adaptive advantages that enhance bacterial survival under selective pressures. To make homologous DNA available in the surrounding environment, *Streptococcus pneumoniae* expresses a cell wall hydrolase (CbpD) that lyses and kills closely related species. This process has been coined *fratricide*, and the acting hydrolase a *fratricin*. A significant fraction of streptococcal species does not encode a CbpD-like protein, but another competence-induced peptidoglycan hydrolase LytF. It has been speculated that LytF serves the same purpose as CbpD, however, our investigations into the role of LytF in *Streptococcus sanguinis* revealed no evidence supporting LytF as a fratricin. Instead, we show that LytF is involved in natural transformation by promoting DNA uptake. An essential part of DNA uptake is the competence-induced type IV pilus, which facilitates DNA uptake by pulling nearby DNA toward the cell. By immunoblotting and microscopy imaging, we found that LytF increases the extracellular levels of the major pilus component ComGC, suggesting that LytF may modify peptidoglycan to promote pilus extrusion across the cell wall, thereby enhancing the efficiency of DNA uptake.

**Importance:** Streptococci are a significant cause of severe infections in both humans and animals. They are particularly adept at acquiring new genes through horizontal gene transfer as they can become competent for natural transformation. This allows them to quickly adapt to selective pressure and spread genes involved in virulence and antibiotic resistance. In *Streptococcus sanguinis*, the competence-induced peptidoglycan hydrolase LytF has been reported to stimulate natural transformation. Our study adds to the understanding of this process by demonstrating that LytF promotes extrusion of the transformation pilus required for DNA uptake.

## Introduction

Most species of the genus *Streptococcus* appear to have the genes required to enter a state in which they become competent for natural genetic transformation (Johnston et al., 2014). During the competent state, streptococci are able to take up naked DNA from the environment and incorporate it into the genome by homologous recombination. The competent state is regulated by the extracellular concentration of specific peptide pheromones (Fontaine et al., 2010; Håvarstein et al., 1995). Two different pheromone-based systems are found to regulate entrance to competence in streptococcal species. The ComCDE system, which was first discovered in *Streptococcus pneumoniae* (Håvarstein et al., 1995; Håvarstein et al., 1996; Pestova et al., 1996), is used by members of the mitis and anginosus phylogenetic groups, including *Streptococcus sanguinis*, while members of the bovis, mutans, salivarius and pyogenic groups use the ComRS system (Fontaine et al., 2015; Shanker et al., 2016). Streptococci expressing the ComCDE system respond to the extracellular concentration of CSP (competence-stimulating peptide encoded by *comC*), whereas species expressing the ComRS system is induced by XIP (*comX*-inducing peptide encoded by *comS*). A large diversity of these two pheromones is found in nature, ensuring species and even strain-specific signalling (Fontaine et al., 2015; Johnsborg et al., 2007; Kilian et al., 2008). When a certain level of extracellular peptide pheromone is reached, it triggers a signalling cascade, which leads to the expression of genes necessary for entering the competent state. The genes transcribed during the competent state are divided into two main categories: the early and late genes. Among others, gene products involved in CSP/XIP production, transport and processing are considered to be early competence genes. The transition from early to late stage of competence is typically defined by the transcription of *comX*, encoding an alternative sigma factor that induces transcription of the late genes (Lee & Morrison, 1999; Peterson et al., 2000). These are genes associated with DNA uptake and homologous recombination (for review see Johnston et al., 2014). In addition, most streptococcal species have a late competence gene encoding a secreted, peptidoglycan hydrolase. There are two main types of these competence-specific peptidoglycan hydrolases: those related to CbpD and those belonging to the LytF group (Berg, Biørnstad, et al., 2012).

CbpD was first described in *Streptococcus pneumoniae* (Guiral et al., 2005; Kausmally et al., 2005). This enzyme is secreted by competent pneumococci and has been shown to attack and lyse non-competent pneumococci as well as members of the closely related species *Streptococcus mitis* and *Streptococcus oralis* (Johnsborg et al., 2008). Because CbpD only kills susceptible sibling cells and close relatives of *S. pneumoniae*, this predatory mechanism has been termed fratricide, and the competence-induced murein hydrolase a fratricin. The fratricide mechanism is proposed as a strategy for the competent cell population to gain access to homologous DNA, which can be integrated into the genomes, thereby improving the chances of obtaining beneficial traits in response to environmental challenges (Johnsborg et al., 2008). To avoid committing suicide by self-produced CbpD, competent pneumococci express an early competence gene encoding an integral membrane protein termed ComM that protects against CbpD (Håvarstein et al., 2006). Evidence suggest that ComM works together with LytR (LytR-CpsA-Psr family protein) to alter the ratio of lipoteichoic to wall teichoic acids (Minhas et al., 2023). In addition, overexpression of ComM results in cell elongation, and it has recently been shown to delay septal peptidoglycan synthesis (Bergé et al., 2017; Juillot et al., 2025; Straume et al., 2017). However, the exact mechanism of ComM-mediated immunity remains unclear. The pneumococcal CbpD consists of an N-terminal <u>c</u>ysteine/<u>h</u>istidine dependent <u>a</u>midohydrolase/<u>p</u>eptidase (CHAP) domain, one or two <u>S</u>rc <u>h</u>omology <u>3b</u> (SH3b) domains and a C-terminal choline binding domain (Cbd) consisting of four choline binding motifs (Layec et al., 2008; Pérez-Dorado et al., 2012). The Cbd module mediates binding of CbpD to choline residues decorating pneumococcal lipoteichoic acid (LTA) and wall teichoic acid (WTA), while the SH3b domain probably binds to the peptidoglycan part of the cell wall (Eldholm et al., 2010). It has been shown that CbpD specifically cleaves nascent peptidoglycan synthesised by the divisome and consequently splits the cell wall of susceptible streptococci along the cell equator (Straume et al., 2020).

As mentioned above, many streptococcal species do not express CbpD-like fratricins during competence. Instead, they express an unrelated peptidoglycan hydrolase termed LytF. Species carrying the *lytF* gene include several members of the mitis and mutans phylogenetic groups, as well as all members of the anginosus and bovis groups. LytF proteins have 3-5 Bsp (group <u>B</u> streptococcal secreted <u>p</u>rotein) repeats at their N-termini and a CHAP domain at their C-terminal end (Berg, Ohnstad, et al., 2012). Muralytic activity has been demonstrated for LytF in *Streptococcus gordonii* and *Streptococcus sanguinis* using zymography, and it has been shown to cause the release of DNA from susceptible oral streptococci (Berg, Ohnstad, et al., 2012; Cullin et al., 2017; Dufour & Lévesque, 2013; Nagasawa et al., 2020; Xu & Kreth, 2013). Localisation studies of LytF in *S. gordonii* showed that the Bsp repeats bind to the septal regions and cell poles, resembling the binding pattern of CbpD (Berg, Ohnstad, et al., 2012). Furthermore, in *S. sanguinis*, the early competence gene *SSA_0195* (new locus tag: *SSA_RS01125*) encodes a protein with weak homology (23.5% identity) to *S. pneumoniae* CbpD-immunity protein ComM (Rodriguez et al., 2011). These results indicate that LytF may be functional analogues to CbpD, but to the best of our knowledge, a fratricide mechanism involving LytF as a lytic enzyme has yet to be described.

In some streptococcal species, mutants lacking LytF or a CbpD-like protein show reduced transformation efficiency when provided similar concentrations of purified DNA, i.e. no fratricide is required to release homologous DNA from target cells (Bjørnstad et al., 2012; Cullin et al., 2017; Eaton & Jacques, 2010; Zhu et al., 2021). Compared to their parental strains, these mutants exhibit reduced transformation rates ranging from 4-fold (*S. sanguinis*) to more than 33-fold (*S. mutans*) (Cullin et al., 2017; Eaton & Jacques, 2010). This puzzling observation has led to speculations that these murein hydrolases may not be involved in fratricide but have a different, or additional, role during competence that influences DNA uptake or recombination (Bjørnstad et al., 2012). An essential structure for DNA uptake is a DNA-binding type IV pilus, which extends from the cell surface to bring the transforming DNA into the DNA uptake machinery. It is encoded by the *comG* operon and, consisting of only five pilins and two assembly proteins, it is the simplest type IV pilus yet discovered. The filament is mainly made up of ComGC major pilin units, while the four minor pilins ComGD-GG form a complex suggested to make up the pilus tip (Mom et al., 2023). Protrusion of the pilus is thought to be powered by the ATPase ComGA docked by a multimer of ComGB constituting the assembly platform (Laurenceau et al., 2013; Pelicic, 2023). Interestingly, the width of the pilus-DNA complex (7-8 nm) is wider than the pore sizes estimated for Gram-positive peptidoglycan (6 nm), suggesting that the cell wall acts as a physical barrier that the pilus needs to penetrate to allow extension into the extracellular milieu (Laurenceau et al., 2013; Pasquina-Lemonche et al., 2020). How this is accomplished by the competent cells is not understood, but it has been hypothesised that specific cell wall remodelling enzymes are required to make space for the transformation pilus, similar to the role such enzymes play in assembling macromolecular structures in Gram-negative bacteria (Koraimann, 2003; Muschiol et al., 2015; Scheurwater & Burrows, 2011).

In the present study, we have investigated the function of LytF in *S. sanguinis* SK36. Our results did not support a role of LytF as a fratricin analogous to the pneumococcal CbpD. Instead, by using non-recombining DNA and immunodetection of the pilus, we show that the reduced transformation efficiency of a Δ*lytF* mutant is caused by a significantly lower extracellular levels of the competence pilus, strongly suggesting that LytF is important for the extrusion of the transformation pilus across the cell wall in *S. sanguinis*.

## Results

### Exploring the potential role of LytF in fratricide

It has been hypothesised that the murein hydrolase LytF is involved in a fratricide mechanism in streptococci, analogous to what has been demonstrated for the murein hydrolase CbpD in *S. pneumoniae*. This is based on the observation that non-competent *S. sanguinis* SK36 treated with LytF results in a growth medium containing more extracellular DNA compared to non-treated cultures, indicating that LytF-mediated cell lysis had taken place (Cullin et al., 2017). If LytF takes part in a fratricide mechanism, it is reasonable to believe that *S. sanguinis* expresses a LytF-immunity protein analogous to the CbpD-immunity protein ComM found in *S. pneumoniae*. In *S. pneumoniae*, deletion of *comM* results in 20 - 30% self-lysis of the competent population and a significant drop in the culture’s optical density due to production of CbpD (Håvarstein et al., 2006; Steinmoen et al., 2002). No LytF immunity protein has been identified, however, *S. sanguinis* SK36 encodes a competence-induced ComM-like protein (*SSA_RS01125*) expressed during the early stage of competence (Rodriguez et al., 2011). We therefore explored whether competence induction could result in self-lysis in Δ*SSA_RS01125* cells, due to exposure to self-produced LytF. Unexpectedly, no decline in OD was observed after induction of competence by addition of exogenous CSP in the Δ*SSA_RS01125* mutant (KP55), suggesting that LytF secreted by the competent KP55 cells did not cause autolysis observable by OD measurements (Figure S1). To further explore the lytic potential of LytF, we performed both live/dead staining and β-galactosidase release assays to detect self-lysis of competent Δ*SSA_RS01125* cells (Fig. 1). The competence-induced Δ*SSA_RS01125* cultures did, however, not show a significant increase in the number of dead cells nor increased release of β-galactosidase, suggesting that LytF is not able to cause self-lysis of Δ*SSA_RS01125* cells (Fig. 1).

**Fig. 1.**
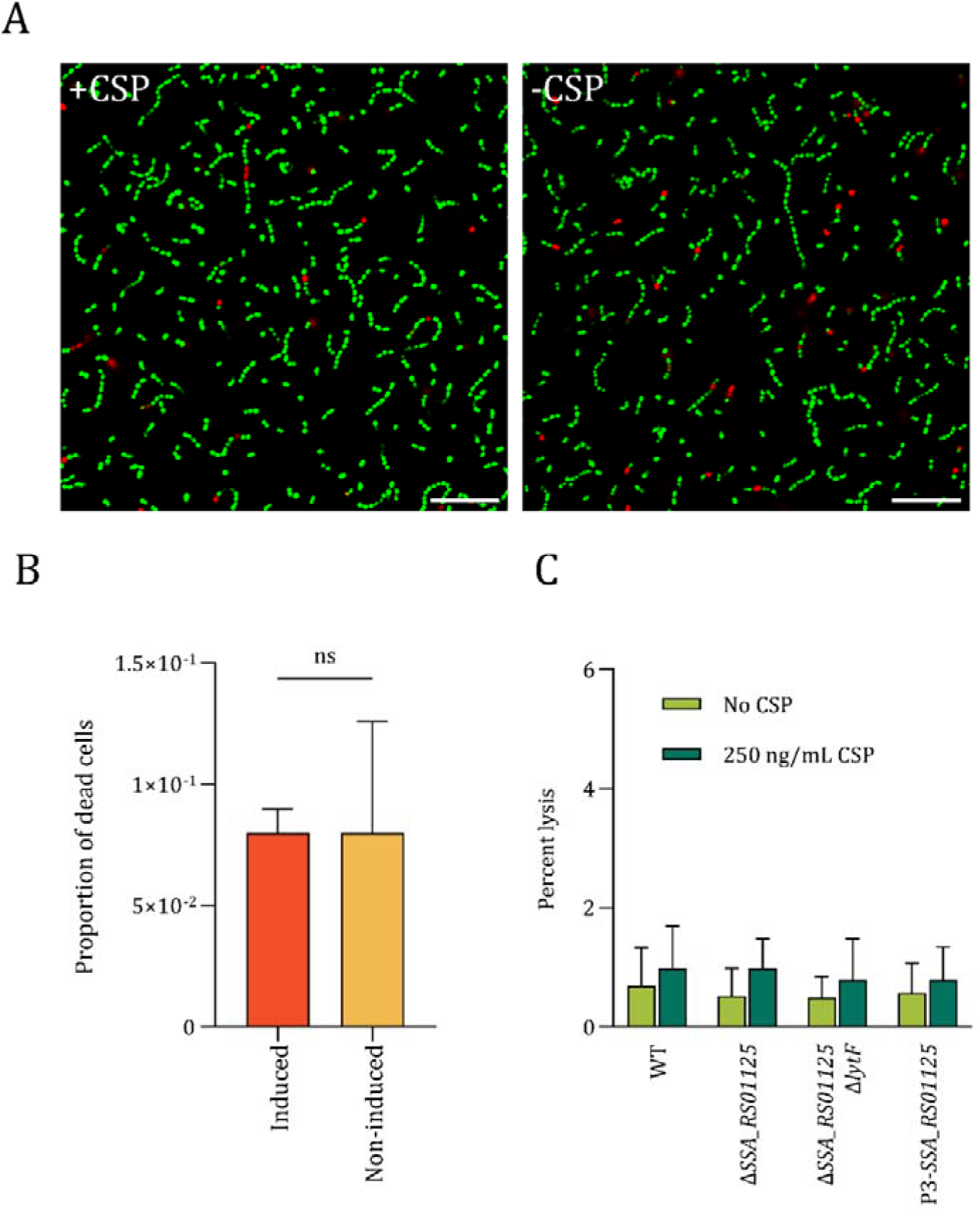
Self-lysis assay of competent Δ*SSA_RS01125* cells. *S. sanguinis* SK36 cells stained using the BacLight Live/Dead kit, which stains live bacteria green and dead bacteria red. Panel A: Live/dead staining of cells induced (+CSP) and non-induced (-CSP) cells of KP55 (Δ*comC*, Δ*SSA_RS01125*). Scale bars are 10 µm. Panel B: Proportion of dead cells in the induced and non-induced cultures. The difference between the means was not statistically significant, as determined by a Student’s *t*-test (p > 0.05). C. Comparison of autolysis in competence-induced *SSA_RS01125*^*+*^ and *SSA_RS01125*□ cells. The following mutant strains were tested: DS921(Δ*comC, lacZ*^+^), DS922 (Δ*comC*, Δ*SSA_RS01125, lacZ*^*+*^), DS934 (Δ*comC*, Δ*SSA_RS01125*, Δ*lytF, lacZ*^+^) and DS926 (Δ*comC*, P3-*SSA_RS01125, lacZ*^+^). In the latter strain, DS926, constitutive expression of SSA_RS01125 is driven by a constitutive synthetic promoter. The four mutants were induced to competence at OD_550_ = 0.3 and grown for another 60 minutes. Non-induced cultures served as negative controls. The relative levels of extracellular β-galactosidase activity suggested that no significant LytF-mediated autolysis occurred in any of the strains during competence.

Since no self-lysis was observed, we wondered whether competent *S. sanguinis* cells become immune to LytF by expressing a different factor than SSA_RS01125. We tested this possibility (i) by measuring cell lysis of non-competent cells treated with purified LytF and (ii) by performing a co-cultivation experiment involving competent attacker cells (LytF producers) and non-competent target cells (Δ*comE* cells) (Fig. 2). To obtain purified LytF we replaced the native *lytF* with a His_6_-version in *S. sanguinis* and purified His_6_-LytF from the supernatant of a competent culture (Fig. S2). His_6_-LytF was then added to exponentially growing Δ*comC* (KP52) cultures at OD_550_ = 0.2. This *comC* knockout mutant is unable to auto-induce competence and, therefore, cannot express any competence-induced immunity factor. Like for the self-lysis experiment, addition of 5 µg/mL His_6_-LytF did not result in cell lysis, as determined by the absence of any decrease in OD_550_ and the lack of fluorescence (SYTOX Green) suggesting that no DNA was released (Fig. 2A).

**Fig. 2.**
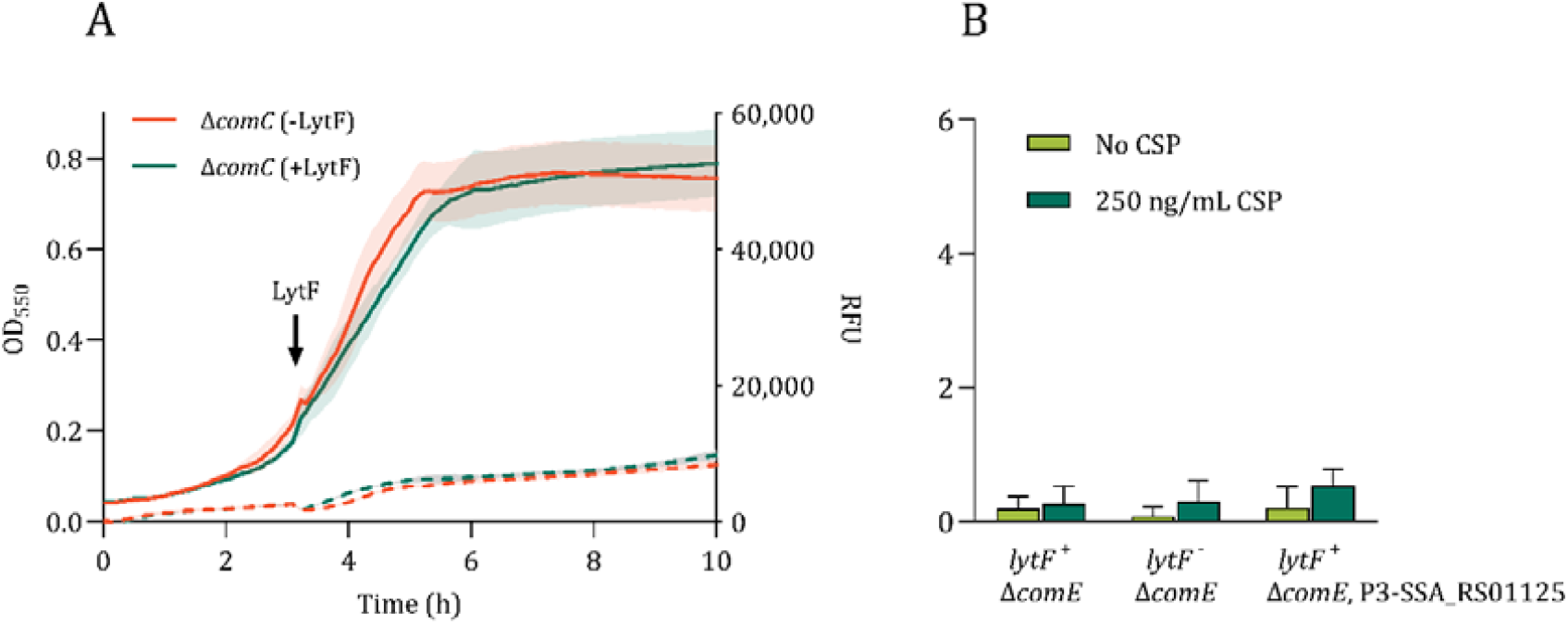
Lytic activity of LytF on non-competent *S. sanguinis* SK36. A. LytF lysis assay using SYTOX. Purified His_6_-LytF was added to a final concentration of 5 µg/mL to a Δ*comC* (KP52) culture at OD_550_ = 0.2. The same volume of PBS was added to a parallel culture as a negative control. The addition of His_6_-LytF did not increase the relative fluorescent signal nor cause a drop in OD_550_, meaning no lysis was detected. B. Percent intracellular β-galactosidase released from non-competent *S. sanguinis* SK36 (Δ*comE*) when co-cultivated with LytF proficient and deficient attacker cells. P3-*SSA_RS01125* cells express SSA_RS01125 constitutively. Attacker cells expressing LytF did not significantly increase the levels of β-galactosidase found in the culture supernatants.

Next, we tested whether intracellularly expressed β-galactosidase was released from non-competent *S. sanguinis* SK36 target cells when grown together with LytF-producing attacker cells. This technique has been successfully used to measure fratricin-mediated lysis in *S. pneumoniae* and *S. thermophilus* (Bjørnstad et al., 2012; Johnsborg et al., 2008; Steinmoen et al., 2002). The *lacZ* gene from *Escherichia coli* was inserted into the middle of a gene of unknown function (*SSA_RS09615*) behind a constitutive extended −10 promoter (see methods) in a *S. sanguinis* Δ*comE* background (unable to induce competence). We set up a series of experiments in which a competence-induced attacker strain secreting LytF was co-cultivated with a competence-deficient (Δ*comE*) target strain. The following pairs of attacker and target strains were tested: KP52 (Δ*comC*) + DS937 (Δ*comE, lacZ*^*+*^), KP61 (Δ*comC*, Δ*lytF*) + DS937 and KP52 + DS939 (Δ*comE*, P3-*SSA_RS01125, lacZ*^+^). However, no significant LytF-mediated release of β-galactosidase was detected in any of the co-cultivation experiments (Fig. 2B). Hence, although LytF has been shown to be a peptidoglycan hydrolase in zymogram assays (Cullin et al., 2017), we could not detect any signs of LytF-induced lysis in competent Δ*SSA_RS01125* or non-competent cells, showing that its lytic activity is limited under the laboratory conditions used here.

### LytF improves DNA uptake in competent *S. sanguinis*

In pneumococci, the fratricide key enzyme (CbpD) has been shown to drastically increase the rate of horizontal gene transfer in a co-culture (Johnsborg et al., 2008), whereas it does not seem to be important for transformation rates when competent cells are provided naked homologous DNA (Fig. S3). For *S. sanguinis, S. mutans, S. thermophilus*, and *S. suis*, on the other hand, their putative fratricins are reported to increase transformation rates when naked DNA is available (Bjørnstad et al., 2012; Cullin et al., 2017; Eaton & Jacques, 2010; Zhu et al., 2021). Since the cells are given equal amounts of homologous DNA, one would reason that it should make the fratricins obsolete in transformation suggesting that the competence-induced murein hydrolases in these species play a role in natural transformation that does not involve lysis of target cells. To investigate how LytF affects the transformation rate, we conducted a series of transformation efficiency assays in relevant mutant strains.

Since transformation efficiency may vary with cell density, we first optimised the transformation protocol by comparing the transformation efficiencies of the wild-type and the Δ*lytF* mutant when induced with CSP at different stages of the exponential growth phase (Table S3). The highest transformation efficiencies were obtained with cultures at OD_550_ = 0.2 - 0.4, resulting in a transformation rate of 0.036-0.054%. Transformation was nearly abolished when inducing at OD_550_ > 0.6. In light of this observation, we performed all transformation experiments at OD_550_ = 0.2. When transforming the KP52 (Δ*comC*) and KP61 (Δ*comC*, Δ*lytF*) strains using a linear DNA Δ*SSA_RS01125::janus* cassette, the Δ*lytF* mutant displayed an average reduction in the number of transformants by approximately 9-fold, which is comparable with the results of Cullin *et al*. (Fig. 3A). The reintroduction of *lytF* with its native promoter in a neutral site (*SSA_RS05615*) in the KP61 genome restored the transformation efficiency to wild-type levels. A strain in which the cysteine (C549) in the active site of the LytF CHAP-domain was substituted for an alanine (RM133) transformed with levels similar to the Δ*lytF*-mutant, demonstrating that the cells require the muralytic activity of LytF for natural transformation to be efficient.

**Fig. 3.**
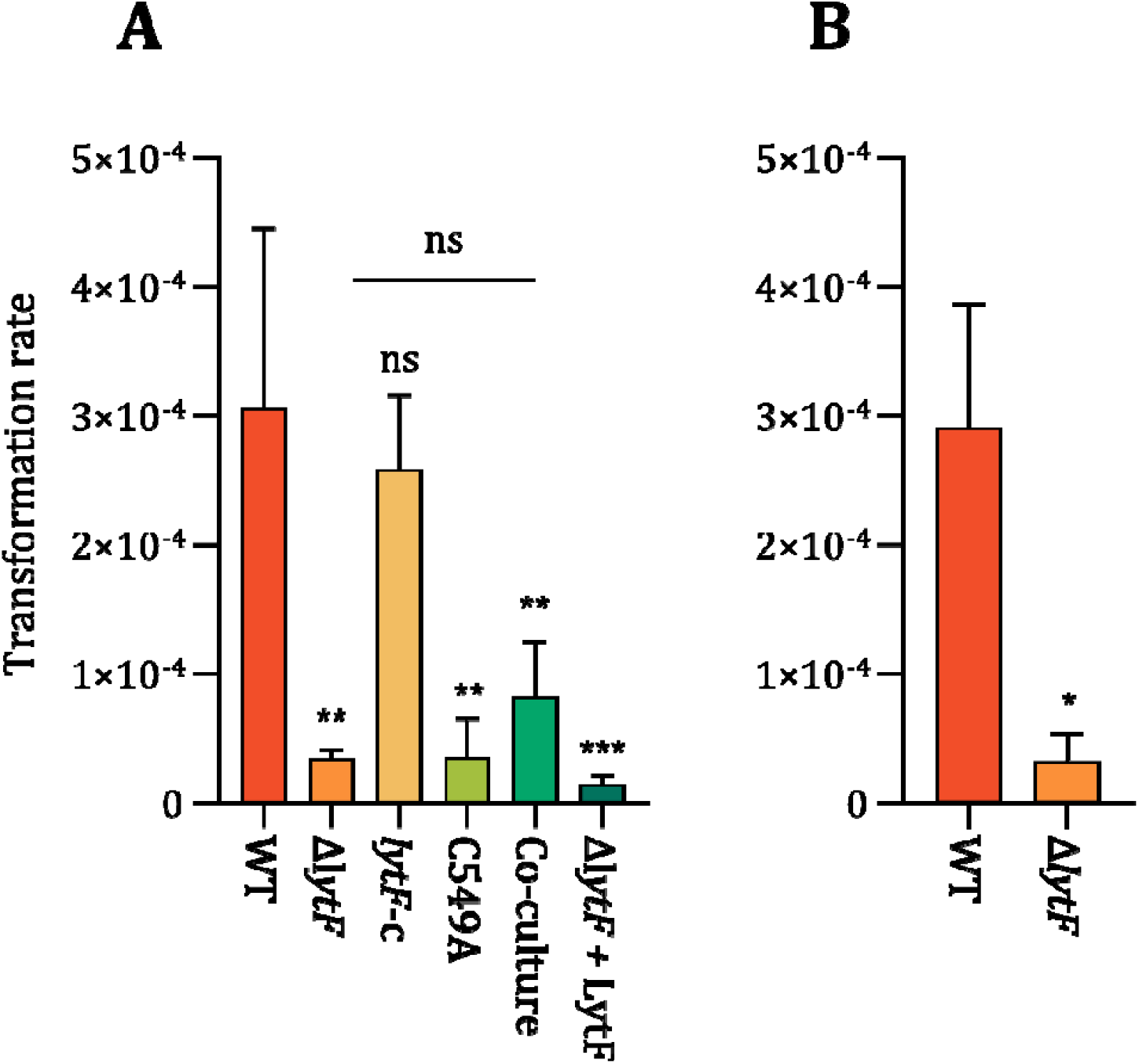
Reduced transformation in *lytF*-deficient cells. Panel A: Cultures were induced to competence at OD_550_ = 0.2, supplied with 250 ng/mL linear transforming DNA (Δ*SSA_RS01125::janus*). The Δ*lytF* mutant (KP61) transformed on average 9 times less efficiently than the wild-type strain (KP52). The transformation rate was restored to wild-type levels in the complementation strain (RM125) in which *lytF* has been ectopically reintroduced into the genome (*lytF-c*). Addition of LytF extracellularly, either by co-cultivation of a *lytF*^-^ strain (KP54) with a *lytF*^+^ strain (KP52), or by supplementing a *lytF*^-^ strain (KP61) with purified His_6_-LytF (+ LytF), did not significantly improve transformation rates. In the co-cultivation experiment, a Δ*SSA_RS01125*::*aad9* cassette was used as the transforming DNA. A strain (RM133) in which the catalytic cysteine in the active site of LytF was substituted for an alanine (C549A) exhibited a transformation rate similar to that of the Δ*lytF* strains. Statistical significance was determined using a one-way ANOVA and Dunnett test, * = p ≤ 0.05, ** = p ≤ 0.01, ns = p > 0.05. Panel B: Same as above, except that the cultures were supplied with transforming DNA in the form of a plasmid (pFD116). In these experiments, the Δ*lytF* mutant transformed on average 11 times less efficiently. Student’s t-test, * = p < 0.05. All experiments were repeated at least three times with similar results.

LytF has a predicted signal sequence for secretion and retains zymogram activity after secretion (Cullin et al., 2017). To determine if the activity of LytF in promoting natural transformation is also maintained extracellularly, we first co-cultured the Δ*lytF*-mutant KP54 (Δ*comC*, Δ*lytF*::*janus*) with the LytF-producing wild-type strain KP52. Co-cultivation did, however, not significantly increase the transformation rate of the mutant (Fig. 3A). In addition, we supplemented the Δ*lytF* mutant strain KP61 with purified His_6_-LytF (5 µg/mL) and, similar to the co-culture results, addition of purified LytF failed to restore the transformation efficiency to wild-type levels (Fig. 3A). This suggests that secreted LytF does not improve transformation rates in strains other than its native producer.

Since natural transformation depends on both DNA uptake and homologous recombination, a disruption in either process could account for the reduced transformation rate seen in fratricin mutants. We hypothesised that the reduced transformation efficiency of the *S. sanguinis* SK36 Δ*lytF* mutant was linked to reduced DNA uptake. To rule out an effect on homologous recombination, we compared transformation rates using linear recombining DNA and a plasmid (pFD116) without sequence homology to the *S. sanguinis* SK36 genome as the donor DNA. Natural transformation with this plasmid only requires its uptake without genomic integration via homologous recombination. Similarly to transformation using homologous DNA, the Δ*lytF* mutant (KP61) transformed on average 11-fold less efficiently than wild-type (KP52) when a plasmid was used as the transforming DNA (Fig. 3B). It thus seems likely that the absence of *lytF* affected the DNA uptake machinery in *S. sanguinis* since the effect on natural transformation persists even when homologous recombination is not involved.

### LytF influences the extracellular amount of the major competence pilin ComGC

An important component of the DNA-uptake machinery in competent streptococci is the type IV competence pilus. In pneumococci it has been shown to bind extracellular DNA and retract, bringing it closer to the cell (Lam et al., 2021; Laurenceau et al., 2013). ComGA and ComGB assemble at the membrane to produce the ComGC pilus filament that crosses the cell wall layer and extends into the extracellular space. To our knowledge, it is not known how the pilus is able to cross the peptidoglycan layer of the cell wall. Bjørnstad, Ohnstad, and Håvarstein (2012) speculated that CbpD of *S. thermophilus* contributes to an increase in transformation efficiency by making modifications to the cell wall that allow the pilus to penetrate the cell surface into the extracellular milieu. Following up on this idea, we compared the amount of the competence pilus of LytF-proficient and -deficient cells. The strains RM32 (Δ*comC*) and RM33 (Δ*comC*, Δ*lytF*) carry a plasmid encoding a C-terminally FLAG-tagged ComGC, which allows for immunodetection of the major pilin. First, we established the dynamics of ComGC expression. Immunoblotting of whole cell extracts of strain RM32 showed that ComGC-FLAG was expressed 10, 20, 30, and 40 minutes after competence induction, with a peak at 30 minutes after induction (Fig. S4). Therefore, samples for comparison of intra- and extracellular pilus levels were taken 30 minutes after competence induction. Notably, immunoblotting of cellular fractions, demonstrated that the amount of ComGC-FLAG in the mutant strain (Δ*lytF*) was dramatically reduced extracellularly and slightly elevated intracellularly compared to the wild-type strain (Fig. 4A). For the complementation strain (*lytF*-c), both extracellular and intracellular fractions were similar to the wild-type (Fig. 4A). The ratio between extracellular and intracellular levels of ComGC was found to be 3-fold higher in the wild-type strain than in the mutant and similar to the complementation strain (Fig. 4B). LytF did not affect production or stability of ComGC as the total amount was comparable across the strains (Fig. S5). These results suggest that the competence pilus relies on the presence of LytF to efficiently extend from the cell during assembly and could explains the Δ*lytF* mutant’s reduced ability for natural transformation.

**Fig. 4.**
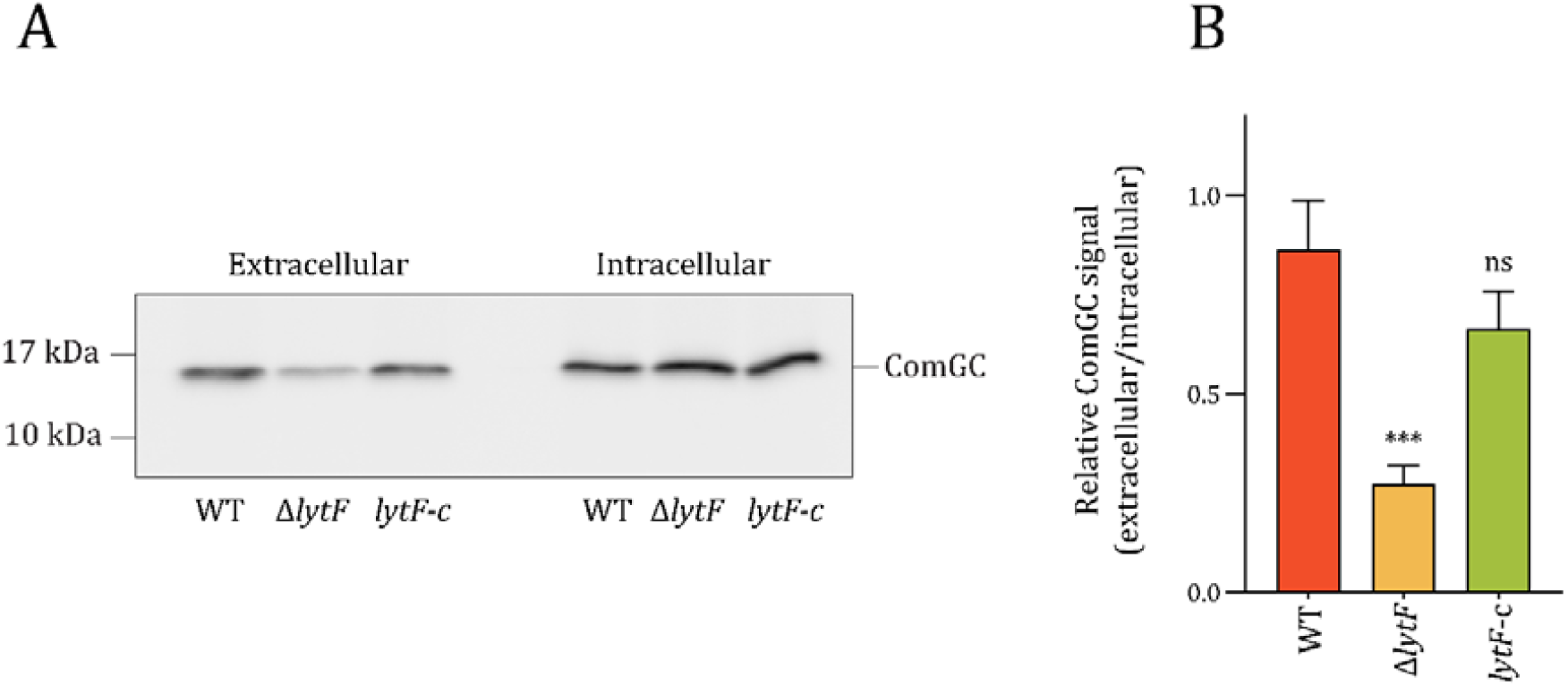
Effect of LytF on intracellular and extracellular levels of ComGC. Panel A: Immunoblots of ComGC in intracellular- and extracellular fractions of the wild-type (RM32), a *lytF*-knockout (RM33), and a *lytF*-complementation strain (RM125). All the strains carry pFD116-P*comG*-*comGC*-*FLAG*. Pili were sheared from competent cells, and the cultures were split into extracellular- and intracellular fractions by centrifugation. Anti-FLAG antibodies were applied to detect ComGC. The experiment was repeated three times with similar results. Panel B: Relative levels of ComGC (extracellular/intracellular) measured from three different immunoblots using ImageJ (Rueden et al., 2017). Statistical significance was determined using a One-way ANOVA and Dunnett’s test, ns = p > 0.05, *** = p ≤ 0.001.

To confirm that the Δ*lytF* mutant indeed had fewer pili exposed on the cell surface, we performed immunofluorescence microscopy (targeting ComGC-FLAG) following the protocol used to visualise pili in *S. pneumoniae* (Laurenceau et al., 2013). In the competence-induced Δ*comC* strain, ComGC-FLAG could be readily detected as fluorescent foci on the cell surface of numerous cells, whereas no foci were seen in the non-competent control cells. Notably, we did not detect any foci in the competence-induced Δ*lytF* strain (Fig. 5). Since we did detect extracellular ComGC-FLAG for the Δ*lytF* mutant using immunoblotting (although significantly less than for wild-type), a total lack of foci was unexpected. Nevertheless, the immunofluorescence results corroborate the observation that fewer pili are found on cells lacking LytF.

**Fig. 5.**
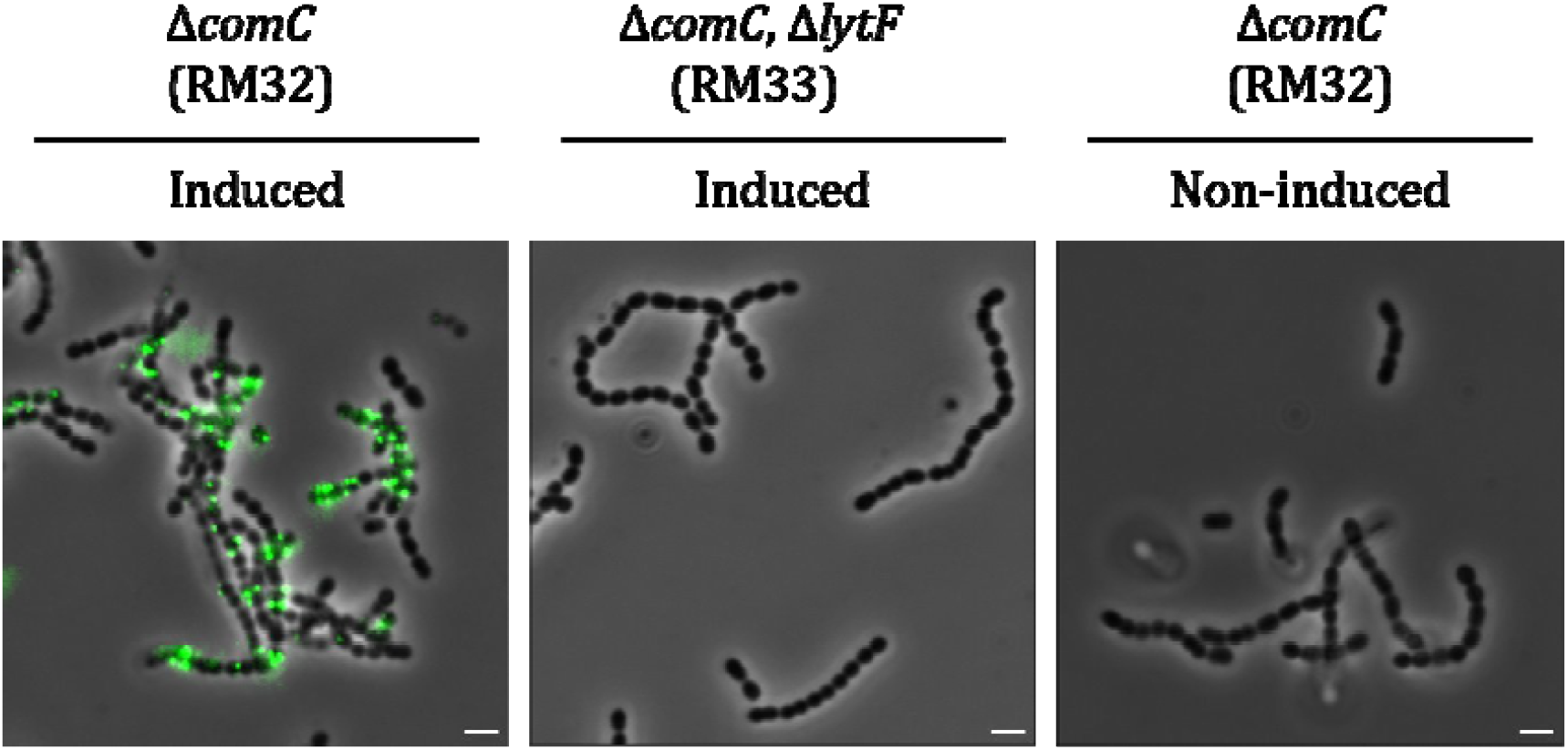
Detection of ComGC using immunofluorescence microscopy. Competent cells of the strains RM32 (Δ*comC*) and RM33 (Δ*comC*, Δ*lytF*), carrying the plasmid pFD116-*comGC*-FLAG, were treated with anti-FLAG antibodies. A non-competence-induced sample of RM32 was included as a negative control. ComGC was detected on induced Δ*comC* cells but was absent from induced Δ*comC* Δ*lytF* cells and the negative control. This experiment was repeated twice with the same results. The images were processed using ImageJ (Rueden et al., 2017). Scale bars are 2 µm.

## Discussion

Competence-induced murein hydrolases in streptococci have previously been shown to be involved in a fratricide mechanism aimed at lysing sibling cells, or cells of closely related species, in order to release homologous DNA that can be taken up by the competent attacker cells and integrated into their genomes by homologous recombination (Håvarstein et al., 2006; Johnsborg et al., 2008; Wei & Håvarstein, 2012). The prime example of this is the pneumococcal CbpD, which is expressed together with the cognate immunity protein ComM. Recently, it was shown that *Streptococcus dysgalactiae* in the pyogenic group also expresses an autolytic murein hydrolase (ScrM) and a cognate immunity protein (ScrI) during competence, representing a system analogous to the pneumococcal CbpD-ComM (Mårli et al., 2025). ScrM displayed high autolytic activity, but it remains to be shown experimentally if ScrM-ScrI is involved in fratricide. For species expressing LytF proteins, on the other hand, no LytF immunity protein has been identified (Berg, Ohnstad, et al., 2012), and it seems to have significantly less capacity to lyse target cells compared to CbpD and ScrM. Still, evidence that LytF is involved in fratricide comes from experiments showing that it increases gene transfer between *S. gordonii* strains and contributes to release of eDNA in *S. gordonii* and *S. mutans* (Berg, Ohnstad, et al., 2012; Nagasawa et al., 2020; Xu & Kreth, 2013). Recently, it has also been shown to contribute to membrane vesicle release in *S. mutans* (Nagasawa et al., 2023). In this work, we explored whether LytF functions as a fratricin in *S. sanguinis* SK36 by testing its ability to lyse non-competent cells as well as a mutant lacking a possible LytF immunity protein (SSA_RS01125, which shares some sequence similarity with the pneumococcal immunity protein ComM). Our observations did not support that *S. sanguinis* LytF functions to lyse sibling cells. Additionally, the lack of autolysis following competence induction of the ΔSSA_RS01125 mutant (Fig. 1, Fig. S1) indicated that this mutant remained immune to self-produced LytF, suggesting that SSA_RS01125 is not an immunity factor. Although SSA_RS01125 shares some sequence similarity to the pneumococcal ComM, *S. sanguinis* SK36 does not encode any ComM homologs (Rodriguez et al., 2011). Besides, as LytF is unrelated to CbpD, it would be reasonable to assume that it requires an immunity mechanism other than pneumococci. However, when we treated non-competent cells with LytF, either as the purified enzyme (Fig. 2) or by an attacker strain (Fig. 3), neither resulted in significant cell lysis compared to competent cells. This suggests that *S. sanguinis* is intrinsically resistant to lysis by LytF and that protection against LytF might not be required. It should be noted, however, that we cannot rule out that LytF contributes to fratricidal efficacy *in vivo* that we were unable to reproduce in the current study.

A striking phenotype observed for some streptococci is that mutants lacking the gene encoding the competence-induced murein hydrolase are less transformable compared to wild type when provided equal amounts of naked DNA, i.e. there is no need for targeted cell lysis to make DNA available for transformation (Bjørnstad et al., 2012; Cullin et al., 2017; Eaton & Jacques, 2010). While deletion of *cbpD* in *S. pneumoniae* R6 (Fig. S3) and *lytF* in *S. gordonii* NCTC 7868 did not significantly reduce the transformability compared to their wild-type counterparts (Berg, Ohnstad, et al., 2012; Johnsborg et al., 2008), a Δ*cbpD* mutant of *S. thermophilus* showed 17-fold reduced transformability (Bjørnstad et al., 2012). More severely, the rate of transformation in a *S. mutans* strain lacking LytF was less than 3% of wild-type levels (Eaton & Jacques, 2010). Cullin *et al*. reported that deletion of the *lytF* gene of *S. sanguinis* SK36 caused a 4-fold reduction in transformability in this bacterium when inducing competence at early exponential phase (Cullin et al., 2017). In line with this, we found that the transformability of the wild-type *lytF*+ strain (KP52) was, on average, 9 times higher than the transformability of the Δ*lytF* mutant (KP61) (Fig. 4A). Interstingly, the same effect was observed when using a plasmid as the transforming DNA (Fig. 4C), which suggests that the DNA-uptake system is somehow affected in the Δ*lytF* mutant. In addition, the transformation rate of a strain harbouring a catalytically inactive version (LytF^C549A^) of the enzyme was comparable to that of the Δ*lytF* mutant (Fig. 4A). This demonstrated that LytF likely facilitates transformation by modifying the cell wall. To our surprise, neither addition of purified His_6_-LytF nor LytF derived from co-culturing with a competent *lytF*+-strain (KP52) restored the transformability of a *lytF*^-^-strain (KP54) (Fig. 3A). This shows that LytF is unable to complement a Δ*lytF* mutant when added from the outside. Ectopic expression of *lytF*, on the other hand, restored transformability to wild-type levels (Fig. 3A). As LytF contains a signal sequence for secretion, these results could mean that LytF is secreted to and is performing its role in the periplasmic space. This could explain the lack of observed lytic activity by extracellular addition.

Streptococcal DNA uptake relies on the type IV competence pili, which act as fishing lines for extracellular DNA (Lam et al., 2021; Laurenceau et al., 2013). During competence, newly synthesised type IV pili alternately extend and retract, presumably functioning as a grappling hook that attaches to extracellular DNA and pulls it across the capsule and peptidoglycan layer. The diameter of the pores of the inner peptidoglycan layer of Gram-positive bacteria has been estimated to be around 6 nm (Pasquina-Lemonche et al., 2020). This is also the estimated diameter of the type IV competence pilus (Laurenceau et al., 2013). However, given that dsDNA is brought into the uptake machinery attaches alongside the pilus, the pores in the peptidoglycan layer need to be at least 8 nm wide, as dsDNA has a width of about 2 nm. Widening of existing pores or creation of pilus-specific pores would therefore be required to make room for the pilus-DNA structure. During assembly of other trans-envelope structures (e.g., bacterial flagella or secretion systems) that are too large to pass through the natural pores of the peptidoglycan sacculus, bacteria are known to produce dedicated peptidoglycan hydrolases which are critical to create the needed space in peptidoglycan (Scheurwater & Burrows, 2011; Vollmer et al., 2008). For example, in *Neisseria gonorrhoeae*, AtlA degrades peptidoglycan, enabling the assembly of a DNA-releasing type IV secretion system without inducing cell lysis (Kohler et al., 2007). Our results established that the *S. sanguinis* Δ*lytF* mutant is less piliated than the wild-type, and complementation of *lytF* from its native promoter restored piliation to wild-type levels (Fig. 4 and 5). We therefore suggest that the Δ*lytF* mutant shows a reduced ability to transform when naked DNA is available because it has fewer extracellular type IV pili to “catch” the DNA. We also showed that normal transformation rates depended specifically on the muralytic activity of LytF (Fig. 3A), suggesting that it aids the competence pilus in entering the extracellular space by making specific cuts in the peptidoglycan layer. The peptidoglycan hydrolase may somehow associate with the pilus and degrade peptidoglycan before or as it extends across the cell wall. Alternatively, LytF may degrade peptidoglycan to enlarge random pre-existing pores, thereby increasing the likelihood of successful pili penetration. Whereas the competence pilus is essential for transformation in *S. sanguinis* (Mom et al., 2023), LytF is not, meaning the pilus extrusion is not entirely dependent on LytF. Our immunoblot displayed that, although to a lesser extent than in the wild-type, ComGC is detected in the extracellular fraction of competent Δ*lytF* cells, suggesting that the pilus is severely hindered but not completely blocked from exiting the cell.

Several streptococcal species encode a LytF-like protein, with the only structural difference between them being the number of Bsp-domains. In *S. mutans*, LytF is an autolytic enzyme that specifically triggers lysis in CSP-responsive cells and is important for eDNA release during biofilm formation (Dufour & Lévesque, 2013; Nagasawa et al., 2020). It has been proposed that this function of LytF is analogous to eukaryotic apoptosis, serving as a response to environmental stress for the overall benefit of the population (Dufour & Lévesque, 2013). But lack of LytF has also been shown to reduce transformability in this species dramatically (Eaton Jacques 2010). In *S. gordonii*, which is more closely related to *S. sanguinis*, LytF strongly increases the rate of gene transfer between two different *S. gordonii* strains during co-cultivation without causing autolysis, but minimally affects the transformation rate in pure cultures (Berg, Ohnstad, et al., 2012; Xu & Kreth, 2013). Our results for LytF from *S. sanguinis* SK36 do not entirely align with those of *S. mutans* LytF or *S. gordonii* LytF, as the former is autolytic while the latter does not affect transformability. Although structurally similar, the LytF hydrolases from different species seem to perform unique roles in the transformation process.

As outlined above, there is data showing that competence-induced peptidoglycan hydrolases in streptococci function as fratricins, facilitators of DNA uptake or both. How can these findings be reconciled? Since solid experimental data support both functions, there seem to be two possible explanations. Either these peptidoglycan hydrolases play different roles in different streptococcal species, or they have a dual function. As no evidence of fratricide was found for *S. sanguinis* LytF, it seems that, at least for this enzyme, the prior hypothesis is more likely. There does not seem to be a clear pattern for predicting which of the two functions the different competence-induced hydrolases perform. Both LytF (*S. sanguinis, S. mutans*) and CbpD-like (*S. thermophilus, S. suis*) enzymes have been reported to affect the transformability, but not all of either variant (Berg, Ohnstad, et al., 2012; Bjørnstad et al., 2012; Cullin et al., 2017; Eaton & Jacques, 2010; Johnsborg et al., 2008; Zhu et al., 2021). The function is also independent of phylogenetic group classification, as the murein hydrolases of *S. pneumoniae* (CbpD), *S. gordonii* (LytF), and *S. sanguinis* (LytF), all belonging to the mitis group, differ in function (Berg, Ohnstad, et al., 2012; Johnsborg et al., 2008).

In this study we could not find evidence supporting LytF to function as a fratricin in *S. sanguinis* SK36, but instead that the expression of LytF influences the transformability of *S. sanguinis* by aiding the competence pilus in entering the extracellular environment.

## Material and Methods

### Bacterial strains and growth conditions

Bacterial strains and species used in this study are listed in Table S1. Todd-Hewitt (TH) broth was used to grow *S. sanguinis* SK36 and its derivatives at 37□C without shaking, except when stated otherwise. When cultivated on TH agar plates, they were grown in anaerobic conditions (<1% O_2_), obtained using AnaeroGen sachets from Oxoid in a sealed container. The *Escherichia coli* strains were grown aerobically in lysogeny broth (LB) or on lysogeny agar (LA) plates at 37□C. When required, TH media were supplemented with 400 µg/mL kanamycin or 200 µg/mL streptomycin. For plasmid retention, 100 µg/mL and 60 µg/mL spectinomycin were used for *S. sanguinis* and *E. coli*, respectively.

### Competence induction and natural transformation

Competence was induced at OD_550_ = 0.2-0.4 using 250 ng/mL CSP (NH2-DLRGVPNPWGWIFGR-COOH). The transforming DNA was added to a final concentration of 250 ng/mL, and the transforming culture was incubated for 2 hours at 37□C before 50-100 µL were plated on TH agar containing the selective antibiotic.

### Construction of mutant strains

Primers used to construct the mutants used in this study are listed in Table S2. The Janus cassette was used to create streptococcal mutants (Sung et al., 2001). The cassette allows for both the selection of its acquisition and its loss. It comprises a kanamycin resistance gene (KmR) and a counter-selectable marker encoding a wild-type streptomycin-sensitive *rpsL* gene (*rpsL*+) that confers dominant streptomycin-sensitivity in a streptomycin-resistant background. Hence, to use the Janus cassette for mutant construction in *S. sanguinis* SK36, we made a streptomycin-resistant mutant of the SK36 strain that contains mutations in its native *rpsL* allele conferring streptomycin resistance. This was achieved by selection on TH agar plates containing 1 mg/mL of streptomycin, giving rise to the streptomycin-resistant mutant KP36.

To delete the *comC* gene of the KP36 strain, the Janus cassette was amplified from genomic DNA obtained from *S. pneumoniae* strain SPH131 using the kp211/kp212 primers. Regions flanking *comC* (approximately 1000 bp) were amplified from genomic DNA of strain SK36 using primers kp207/kp208 and kp209/kp210. The three PCR products were fused by overlap extension PCR using primers kp207/kp210 (Higuchi et al., 1988). The resulting amplicon was introduced into KP36 through natural transformation. The Janus cassette was subsequently removed by transformation with a PCR construct consisting of only the fused flanking regions, followed by selection on streptomycin-containing TH-agar plates. This resulted in the mutant KP52, in which only the coding region of the *comC* gene had been excised. All genetic modifications introduced to *S. sanguinis* were performed using this approach. Strains RM88, RM104, and RM133 were created by removing the Janus cassette, replacing *lytF* with PCR constructs comprised of the *lytF*-flanking regions fused to the modified variants of the gene (*lytF-sfGFP, his-lytF*, and *lytF*_C549A,_ respectively). Transformants were verified by colony PCR and Sanger sequencing (GATC, Eurofins).

For β-galactosidase assays, *lacZ* was introduced to *S. sanguinis* by placing it in the *hirL* locus (*SSA_RS09615*) similarly to the approach used for *S. pneumoniae* (Steinmoen et al., 2002). A Δ*hirL*::janus cassette was constructed by amplifying janus using primers Kan484F/RpsL41R and gDNA from RH426 as template. The *hirL* upstream- and downstream regions were amplified from genomic DNA of *S. sanguinis* SK36 using primer pairs ds703/ds704 and ds705/ds706, respectively. Then the three amplicons were fused by using primer pair ds703/ds706. A Δjanus::P3-*lacZ* amplicon was used to replace janus in the *hirL* locus with P3-*lacZ* (transcription of *lacZ* is driven by a constitutive synthetic P3 promoter with sequence 5’-TTGCACTGTCCCCCTGGTATAATAACTATACATGCAAGATCTAAAT-3’). P3-*lacZ* was amplified from genomic DNA derived from strain RH2 using primers ds711 and ds712. For replacement of janus the P3-*lacZ* amplicon was fused to the *hirL* upstream- and downstream region. The upstream region was amplified using primer pair ds703/ds713 and the downstream region using primers ds706/ds714 and *S. sanguinis* SK36 gDNA as template. Finally, primers ds703 and ds706 were used to fuse the three amplicons and produce Δjanus::P3-*lacZ*.

The pFD116-P*comGA*-*comGC*-FLAG plasmid was constructed by first amplifying the promoter region (156 bp upstream *comGA*) of the *comG*-operon and the *comGC* gene from *S. sanguinis* SK36. Primer pairs rm032/rm035 and rm033/rm034 were used, respectively. Primer rm034 contains a FLAG tag-encoding sequence. The amplified fragments were joined by overlap extension PCR, cleaved with SpeI and NcoI, and ligated into pFD116. Next, the ligation reaction was transformed into chemically competent *E. coli* cells by heat shock at 42□C. Finally, the plasmid sequence was verified by Sanger sequencing (GATC, Eurofins) and transformed into *S. sanguinis* by natural transformation, as described above.

### Purification of His_6_-LytF

His_6_-LytF was purified from the supernatant of a competent *S. sanguinis* culture (RM104), carrying the *lytF* gene fused with 6x-*his*, using immobilised metal affinity chromatography (IMAC). A 500 mL culture of OD_550_ = 0.2 was induced to competence and incubated at 37□C for 1 hour. After centrifugation at 5,000 × g for 10 minutes, the supernatant containing His_6_-LytF was filtered using a 0.45 µm filter. The filtrate was then passed through a 1 mL HisTrap^TM^ HP column (Cytiva Life Sciences) with the Äktaprime plus system (GE Healthcare). The column was washed with 10 column volumes of 10 mM Tris-HCl (pH 7.4), 150 mM NaCl, 20 mM imidazole before His_6_-LytF was eluted using a linear gradient of imidazole from 20-500 mM in the same buffer conditions. The eluate was dialysed with a buffer of 10 mM Tris-HCl, 150 mM NaCl, pH = 7.4, for 1 hr at room temperature. The presence and purity of the protein were assessed by SDS-PAGE electrophoresis (Fig. S1) before it was stored at −80□C.

### β-galactosidase assay

Exponentially growing *S. sanguinis* SK36 mutants expressing *lacZ* constitutively were diluted to OD_550_ = 0.05 in a final volume of 5 mL TH broth. The bacteria were grown to OD_550_ = 0.3 and induced to competence. Uninduced cultures were grown in parallel as a negative control. For co-cultivation experiments, 2.5 mL of the attacker and 2.5 mL of the target cells were mixed at OD_550_ = 0.3, and CSP was added to induce competence in the attacker cells. After incubation at 37°C for 60 minutes, the β-galactosidase activity released to the growth medium and total β-galactosidase activity in the culture (growth medium and intracellular) was quantified. Three mL from each cell culture was used as follows: one mL was sterile filtered through a 0.2 µm filter to obtain cell-free culture supernatants, one mL cell culture was lysed using bead beating with 0.5 g ≤ 106 µm glass beads (Sigma) for 3 × 20 seconds at 6.5 m/s, and one mL was used to measure OD_550_. The β-galactosidase activity was detected by adding 960 µL culture supernatant or 200 µL clarified cell lysate to 240 µL 5x Z-buffer (5 mM MgCl_2_, 250 mM β-mercaptoethanol, 50 mM KCl, 0.3 M Na_2_HPO_4_, 0.2 M NaH_2_PO_4_) containing 4 mg/mL o-nitrophenyl-β-D-galactopyranoside (ONPG). For the cell lysate samples, 760 µL TH medium was added to a final volume of 1200 µL. The samples were incubated at 37 °C for 90 minutes before the reaction was stopped by adding 500 µL 1 M Na_2_CO_3_. ONPG is hydrolysed by β-galactosidase to galactose and ortho-nitrophenol; the absorption of the latter was measured at 420 nm. The β-galactosidase activity was calculated according to Miller *et al*. (1972).

### SYTOX Green nucleic acid stain assay

The Δ*comC* mutant (KP52) was grown to OD_550_ ∼ 0.2 in C-medium (Lacks & Hotchkiss, 1960) before being diluted to OD_550_ = 0.05. Here, C-medium was used instead of TH as its lack of colour ensures minimal interference with the fluorescent signal. The culture was transferred to a transparent bottom black polystyrene 96-well plate (Merck), and SYTOX^TM^ Green Nucleic Acid stain (ThermoFisher) was added to a final concentration of 2 µM. The stain fluoresces when binding nucleic acids but is unable to cross intact cell membranes. Optical density (550 nm) and fluorescence (excitation at 485 nm and emission at 535 nm) were measured using the HIDEX Sense microplate reader. At OD_550_ = 0.2, a final concentration of 5 µg/mL His-LytF was added to the culture.

### Transformation efficiency assays

The transformation efficiency assays were performed by natural transformation using a Δ*SSA_RS01125::*Janus cassette. Following transformation, the cultures were serially diluted and plated on TH plates with and without kanamycin. The plates were incubated anaerobically overnight, and colonies were counted the next day. Colonies on the antibiotic- and non-antibiotic plates were used to estimate transformant- and total cell count, respectively. The transformation efficiency was calculated as the number of transformants divided by the total cell count and was based on at least three repetitions of each experiment. Only plates containing between 30 and 300 colonies were included in the calculations. To assess at what OD_550_ *S. sanguinis* SK36 transformed the most efficiently, the assay was performed as described above, except samples of a growing culture were taken for every timepoint, representing various OD_550_ values ranging from 0.05 to 0.9 with increments of 0.1. For the co-cultivation assay, a culture consisting of KP52 and KP54 in equal amounts was prepared in parallel right before competence induction and transformed using a Δ*SSA_RS01125::aad9* cassette.

### Western blotting

Competence was induced in a 5 mL culture at OD_550_ = 0.2, which was then incubated for 30 minutes at 37□C. The pilus appendages were mechanically detached from cells by vortexing for the last 10 minutes of incubation. Cells and supernatant were split by centrifugation at 4,000 × g for 10 minutes. After decanting the supernatant containing the sheared pili, the cells were washed once with PBS and lysed in 200 µL PBS using the FastPrep®-24 homogeniser (MP Bio). All samples were normalised based on total protein amounts of the whole cell extracts dissolved in 1% SDS estimated by measuring the A280 nm. Following SDS-PAGE (15% resolving gel) and transfer of the separated proteins onto a methanol-activated PVDF membrane using electroblotting, the membrane was blocked in 5% skim milk in TBS-T (w/v) at room temperature for 1 hour and then at 4°C O/N. The membrane was incubated with anti-FLAG antibodies (1:4000, ThermoFisher) and anti-rabbit antibodies (1:4000, HRP, ThermoFisher) for 1 hour each and was washed with TBS-T (3 × 10 min) after each incubation. Finally, secondary antibodies associated to the membrane was revealed using the SuperSignal^TM^ West Pico PLUS Chemiluminescent Substrate (ThermoFisher) and the Azure c400 Imaging System (Azure Biosystems).

For the time-series experiment, cultures were incubated for 10-40 minutes with increments of 10 following competence induction. A non-induced culture was included as a negative control (time 0). SDS-PAGE and blotting were performed as described above.

### Microscopy

Fluorescence microscopy for visual detection of ComGC was performed as outlined by Laurenceau *et al*. (2013) with minor modifications. Briefly, the cells were induced to competence and incubated for 30 minutes. They were then harvested and resuspended in 500 µL PBS. A 50 µL aliquot of this suspension was allowed to adhere to a poly-L-lysine slide for 5 minutes. This and all the following steps were conducted in a humidity chamber. The cells were fixed with 3.7% formaldehyde in PBS, blocked with 1% BSA for 5 minutes, then incubated with anti-FLAG antibodies (1:4000, ThermoFisher) for 1 hour, followed by an incubation with anti-rabbit antibodies (1:4000, Alexa Fluor™ 488, ThermoFisher) for another hour. In between these steps, the cells were washed with PBS. Images were taken with a Zeiss LSM700 microscope and analysed using ImageJ (Rueden et al., 2017).

For live/dead imaging, the BacLight Live/Dead staining kit (Invitrogen) was used to stain planktonic cultures as described by the manufacturer. Images were taken using a 555-nm laser line for the excitation of propidium iodide, and a 488-nm laser line was applied for the excitation of SYTO 9. Zen software (ZEN Zeiss Lite) and the ImageJ plugin MicrobeJ was used for image analysis (Ducret et al., 2016; Rueden et al., 2017).

## Acknowledgements

We thank Zhian Salehian for technical assistance with cloning. The project was supported by a grant from the Norwegian University of Life Sciences.

