## Supplementary material for "LytF contributes to pilus extrusion during natural competence in *Streptococcus sanguinis* SK36": Supplemetal material

**Supplemental tables**

**Table S1.** Bacterial strains used in this study.

| **Strain** | **Relevant characteristics** | **Source or reference** |
| --- | --- | --- |
| *S. pneumoniae* strains | | |
| R704 | R6 derivative, *comA*::*ermAM* | J. P. Claverys^a^ |
| RH425 | R704, but streptomycin resistant | (Johnsborg & Håvarstein, 2009) |
| RH426 | RH425, contains the janus cassette downstream of *amiF* | (Johnsborg & Håvarstein, 2009) |
| RH2 | R704 having *lacZ* integrated into *hirL* | (Johnsborg et al., 2008) |
| RM17 | RH425, but Δ*cbpD* | This study |
| MG12 | Clinical strain from NIPH, Δ*cps*::*aad9* |  |
| RM30 | MG12, but Δ*cbpD* | This study |
| *S. sanguinis* SK36 strains | | |
| KP36 | ATCC BAA-1455 | This study |
| KP52 | Δ*comC* | This study |
| KP55 | Δ*comC*, Δ*SSA_RS01125* | This study |
| KP61 | Δ*comC*, Δ*lytF* | This study |
| KP66 | Δ*comC*, Δ*lytF*, Δ*SSA_RS01125* | This study |
| DS921 | Δ*comC*, *lacZ*+ | This study |
| DS922 | Δ*comC*, Δ*SSA_RS01125*, *lacZ*+ | This study |
| DS926 | Δ*comC*, P3-*SSA_RS0112*5, *lacZ+* | This study |
| DS934 | Δ*comC*, Δ*SSA_RS01125*, Δ*lytF*, *lacZ*+ | This study |
| DS937 | Δ*comE*, *lacZ*+ | This study |
| DS939 | Δ*comC*, Δ*comE*, P3-*SSA_RS0112*5, *lacZ*+ | This study |
| RM32 | Δ*comC*, pFD116-P*comGA*-*comGC*-*FLAG* | This study |
| RM33 | Δ*comC*, Δ*lytF*, pFD116-P*comGA*-*comGC*-*FLAG* | This study |
| RM88 | Δ*comC*, Δ*lytF-CHAP*::*lytF-sfgfp* | This study |
| RM104 | Δ*comC*, Δ*lytF*::6x*his-lytF* | This study |
| RM125 | Δ*comC*, Δ*lytF*, ΔSSA_RS05615::*lytF*-*ermB*, pFD116-P*comGA*-*comGC*-*FLAG* | This study |
| RM133 | Δ*comC*, Δ*lytF::lytF_C549A_* |  |
| *E. coli* strains | | |
| RM18 | DH5α containing pFD116-P*comG*-*comGC*-*FLAG* | This study |

^a^ Gift from J.P. Claverys

**Table S2.** Primers used in this study.

| **Primer** | **Sequence (5’-3’)** | **Application** | **Source** |
| --- | --- | --- | --- |
| Kan484F | GTTTGATTTTTAATGGATAATGTG | Janus cassette | Johnsborg 2008 |
| RpsL41R | CTTTCCTTATGCTTTTGGAC |  |  |
| kp171 | ATCAGTATTAGTTCCTGACTCG | Δ*SSA_RS01125*::Janus::DEL | This study |
| kp172 | TGAACCTCCAATAATAAATATTCTCTCCATTCTTCTCTTATC |  |  |
| kp173 | TTTCTAATATGTAACTCTTCCCAATAAAGTGTTTTGTCAAA  TAGAAGAAT |  |  |
| kp174 | CCTGAGTCTCAGGATTGACC |  |  |
| kp175 | GATAAGAGAAGAATGGAGAGAATATTTATTATTGGAGGTTCA |  |  |
| kp176 | ATTCTTCTATTTGACAAAACACTTTATTGGGAAGAGTTACAT  ATTAGAAA |  |  |
| kp177 | GATAAGAGAAGAATGGAGAGAATAAAGTGTTTTGTCAAATA  GAAGAAT |  |  |
| kp178 | ATTCTTCTATTTGACAAAACACTTTATTCTCTCCATTCTTCTCT  TATC |  |  |
| kp179 | ACTGGACTATTCCATCGC | Δ*lytF*::  Janus::DEL |  |
| kp180 | GCCATCCAGGAACTCCTCG |  |  |
| kp181 | GTCCAAAAGCATAAGGAAAGTTTCTAATATGTAACTCTTCCC  AATAAAACTCCTTTGTGAGAATGG |  |  |
| kp182 | CACATTATCCATTAAAAATCAAACTGAACCTCCAATAATAAA  TGTCGCTTTCGCACTTCTC |  |  |
| kp183 | GAGAAGTGCGAAAGCGACAAAACTCCTTTGTGAGAATGG |  |  |
| kp184 | CCATTCTCACAAAGGAGTTTTGTCGCTTTCGCACTTCTC |  |  |
| kp193 | CGACCTTATTAGCGGTATACC | *lytF* |  |
| kp194 | GACCTTCAGTGGTATGACG |  |  |
| kp195 | TGGCTATGTTGACTCAGACG | *SSA_RS01125* |  |
| kp196 | AATCATATTTCTGAGAATATGGG |  |  |
| kp207 | TTATATGAAGTAGTTACACTGG | Δ*comC*::Janus::DEL |  |
| kp208 | TGAACCTCCAATAATAAATAACTATCTCCTATCTTTTTATCTTG |  |  |
| kp209 | TTTCTAATATGTAACTCTTCCCAATGAATTTTAAAGTTTAGGG  GAGA |  |  |
| kp210 | AGCTGAATCTGATGTTCACG |  |  |
| kp211 | CAAGATAAAAAGATAGGAGATAGTTATTTATTATTGGAGGTT  CA |  |  |
| kp212 | TCTCCCCTAAACTTTAAAATTCATTGGGAAGAGTTACATATT  AGAAA |  |  |
| kp213 | CTATTCTTTTCAAATTGCTTTAAATAGC |  |  |
| kp214 | GCTATTTAAAGCAATTTGAAAAGAATAG |  |  |
| kp215 | ATTCGCCTTCTAAGCGAACG |  |  |
| kp216 | CGGAACTATAATCTCAAGACC |  |  |
| kp217 | AATCCAATTATTTCAAGTGACATAACTATCTCCTATCTTTTTA  TCTTG |  |  |
| kp218 | CAAGATAAAAAGATAGGAGATAGTTATGTCACTTGAAATAA  TTGGATT |  |  |
| kp219 | CATGATGGATACTACAGTGC |  |  |
| kp220 | CTACTAATCATCAGAATCTGAG |  |  |
| ds703 | TGTCCAAGCTGGTCAAGGAAC | Δ*hirL*::Janus |  |
| ds704 | CACATTATCCATTAAAAATCAAACGTTCAAGCCTCCTTGATT  CAC |  |  |
| ds705 | GTCCAAAAGCATAAGGAAAGAGAAAATTCAGAATTATTTAA  TTTGTTC |  |  |
| ds706 | TCCTGGACGAGAATTACGAC |  |  |
| ds711 | TTGCACTGTCCCCCTGGTATAATAACTATACATGCAAGATCTA  AATAGGAGGAAAATTAGTGGAAGTTACTGACGTAAGATTAC | Δ*hirL*::Janus::P3-*lacZ* |  |
| ds712 | TTATTTTTGACACCAGACCAAC |  |  |
| ds713 | TACCAGGGGGACAGTGCAAGTTCAAGCCTCCTTGATTCAC |  |  |
| ds714 | GTTGGTCTGGTGTCAAAAATAAAGAAAATTCAGAATTATT  TAATTTGTTC |  |  |
| ds720 | GGCTTGTTTCTGACATGTATC | Δ*comE*::Janus |  |
| ds721 | CACATTATCCATTAAAAATCAAACAAATTCTATCTCCTAATTG  TTAAAATC |  |  |
| ds722 | GTCCAAAAGCATAAGGAAAGAAACCTGATATAATGGAATATG  TTC |  |  |
| ds723 | AAGTAGATGCTATAACTACTAATG |  |  |
| rm032 | GATCACTAGTAAACTAAAAACCACTAAGCCTCTTT | pFD116-P*comG*-*comGC*-*FLAG* |  |
| rm033 | ATGAAAAAATTTAATACCTTAAAAGTTCA |  |  |
| rm034 | CATGCCATGGTTACTTGTCGTCATCGTCTTTGTAGTCATTGGCAACCGCTTGA |  |  |
| rm035 | TGAACTTTTAAGGTATTAAATTTTTTCATATATCCTCCTCACCTTACTATTCG |  |  |
| rm120 | TGTATTGGGTCCCTTGTCGC | *lytF-CHAP*::*lytF-sfGFP* |  |
| rm121 | CACCTTTAGAACCGGTAAGATGTTTACCACCACCACCACCCTCAACCTGATAGCTAGTTCCAGTC |  |  |
| rm122 | GACTGGAACTAGCTATCAGGTTGAGGGTGGTGGTGGTGGTAAACATCTTACCGGTTCTAAAGGTG |  |  |
| rm123 | GAGAAGTGCGAAAGCGACTTATTATGCGGCCGCTCCACTAG |  |  |
| rm124 | CTAGTGGAGCGGCCGCATAATAAGTCGCTTTCGCACTTCTC |  |  |
| rm125 | ATCAGACGGCCTTGCCACTG |  |  |
| rm129 | CACATTATCCATTAAAAATCAAACAAAACTCCTTTGTGAGAATGGTTTTGG |  |  |
| rm130 | CCAAAACCATTCTCACAAAGGAGTTTTGTTTGATTTTTAATGGATAATGTG |  |  |
| rm131 | CGTCCAAAAGCATAAGGAAAGGTCGCTTTCGCACTTCTCAAAACTAC |  |  |
| rm132 | GTAGTTTTGAGAAGTGCGAAAGCGACCTTTCCTTATGCTTTTGGACG |  |  |
| rm152 | TTGAGAAGTGCGAAAGCGACTTATTTAGGGTAGATATAGGTCAGGC | *6x-his*-*lytF* |  |
| rm153 | GTCGCTTTCGCACTTCTCAA |  |  |
| rm154 | ATGATGATGATGATGATGCGCATGGACTGTTGAGTGG |  |  |
| rm155 | CATCATCATCATCATCATACCGAAACGCCTGAATCTG |  |  |
| ds121 | TTATAATTTTTTTAATCTGTTATTTAAATAG | Δ*S*SA_RS01125::*aad9* |  |
| ds717 | TTAAATGTGCTATAATACTAGAAAATACTTGTGGAGGTTCCATTGTGAGGAGGATATATTTGAATAC |  |  |
| rm161 | CTATTTAAATAACAGATTAAAAAAATTATAAAAAGTGTTTTGTCAAATAGAAGAATTATGG |  |  |
| rm162 | TCCACAAGTATTTTCTAGTATTATAGCACATTTAAATTCTCTCCATTCTTCTCTTATCATTATAGAAGAAG |  |  |
| rm203 | ACTCAAGGATACAACGATTGTCTCAGC | ΔSSA_RS05615::*lytF*-*ermB* |  |
| rm204 | ACGTCCAAAAGCATAAGGAAAGTTAAAAACACCCCAAAAGTTAGATTTTTTCTG |  |  |
| rm205 | GATTATATCACATTATCCATTAAAAATCAAACTTATTTGTCTAACTTTTTGGGGTCAGTACAC |  |  |
| rm206 | GCTCAGGCCCTGCTAAGTCG |  |  |
| rm207 | TTATTTAGGGTAGATATAGGTCAGGCG |  |  |
| rm208 | TCCATTCTAATGAGAAAAAGAATATCTCTTCTATATGATCG |  |  |
| rm209 | CGCCTGACCTATATCTACCCTAAATAATTAAAAACACCCCAAAAGTTAGATTTTTTCTG |  |  |
| rm210 | ATCATATAGAAGAGATATTCTTTTTCTCATTAGAATGGATTATTTGTCTAACTTTTTGGGGTCAGTACAC |  |  |
| rm270 | GGTCAAGCTACTTGGGGAGC | Δ*lytF*::*lytF*_C549A_ |  |
| rm271 | CTCCCCAAGTAGCTTGACCG |  |  |

**Table S3.** Transformation efficiency of *S. sanguinis* mutants at different ODs during exponential growth phase.

|  | KP52 (Δ*comC*) | KP61(Δ*comC*, Δ*lytF*) | KP55 (Δ*comC*, Δ*S*SA_RS01125) | KP66 (Δ*comC*, Δ*S*SA_RS01125, Δ*lytF*) |
| --- | --- | --- | --- | --- |
| OD_550_ | Transformation  efficiency (%)* | Transformation  efficiency (%) | Transformation  efficiency (%) | Transformation  efficiency (%) |
| 0.05 | 0.007 | 0.003 | 0.017 | 0.0016 |
| 0.1 | 0.018 | 0.002 | 0.01 | 0.0007 |
| 0.2 | 0.04 | 0.0014 | 0.048 | 0.0022 |
| 0.3 | 0.036 | 0.0035 | 0.029 | 0.0017 |
| 0.4 | 0.054 | 0.0033 | 0.055 | 0.0024 |
| 0.5 | 0.012 | 0.0011 | 0.011 | 0.0008 |
| 0.6 | 0.002 | 0.0001 | 0.0033 | 0.0001 |
| 0.7 | 0.0001 | 0 | 0 | 0 |
| 0.8 | 0 | 0 | 0 | 0 |
| 0.9 | 0 | 0 | 0 | 0 |

* Calculated as the number of CFUs of transformants/total number of CFUs in the culture.

**Supplemental figures**

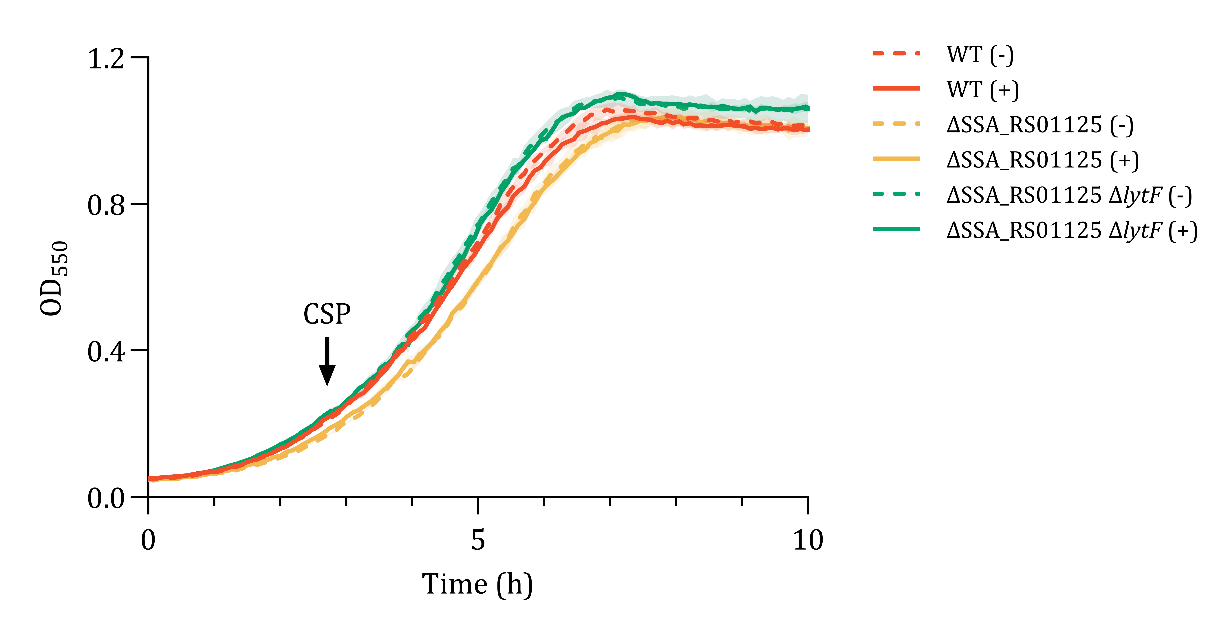

**Fig. S1** Growth curve assay of competent (+) and non-competent (-) *S. sanguinis* WT (KP52), Δ*SSA_RS01125* (KP55), and Δ*SSA_RS01125*, Δ*lytF* (kP66). At OD_550_ = 0.2, the (+)-cultures were induced to competence by the addition of CSP (final conc. 250 ng/mL) and the (-)-cultures were supplied with the same volume of PBS. The experiment was repeated three times with similar results.

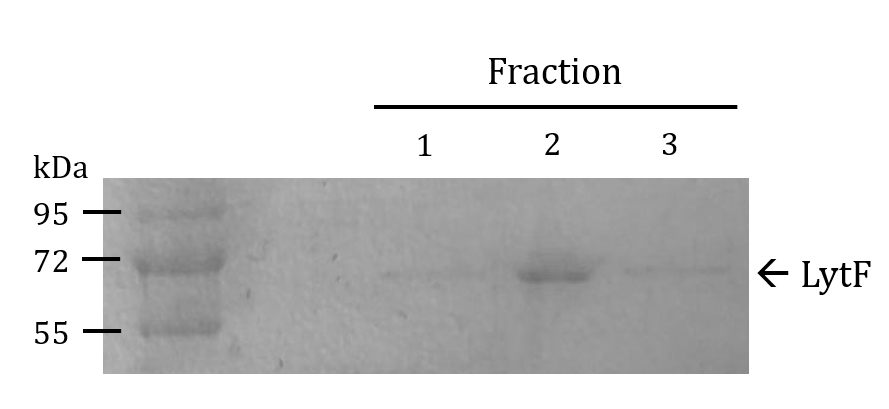

**Fig. S2** SDS-PAGE of immobilised metal affinity chromatography (IMAC) purified His_6_-LytF. The histidine-encoding sequence was inserted just downstream of the secretion signal sequence in *lytF*. The protein was expressed natively and purified form the supernatant of the competent culture. The numbers 1, 2, and 3 correspond to elution fractions with the highest A280 nm absorbance during imidazole elution from the Ni-NTA column. A protein band corresponding to the size of His_6_-LytF (73 kDa) was visualised by Coomassie brilliant blue staining.

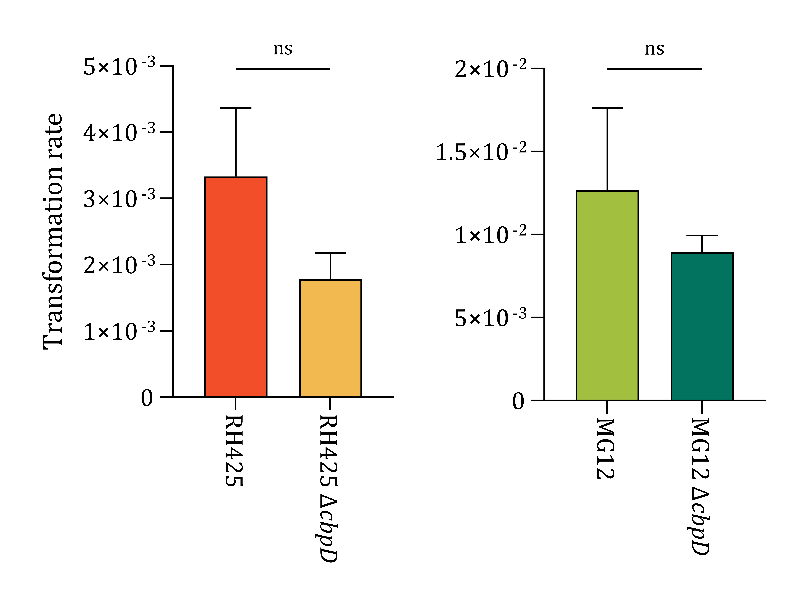

**Fig. S3** Transformation rate of *S. pneumoniae* Δ*cbpD*-mutants of a lab strain (RH425) and a clinical isolate (MG12) compared to their parental strains. The transformation rate was calculated as the number of CFUs of transformants divided bythe total number of CFUs in the culture. The difference in transformation rates between the Δ*cbpD*-mutant and the parental strain was non-significant (Student’s t-test, p = 0.2666). The experiment was repeated three times with similar results.

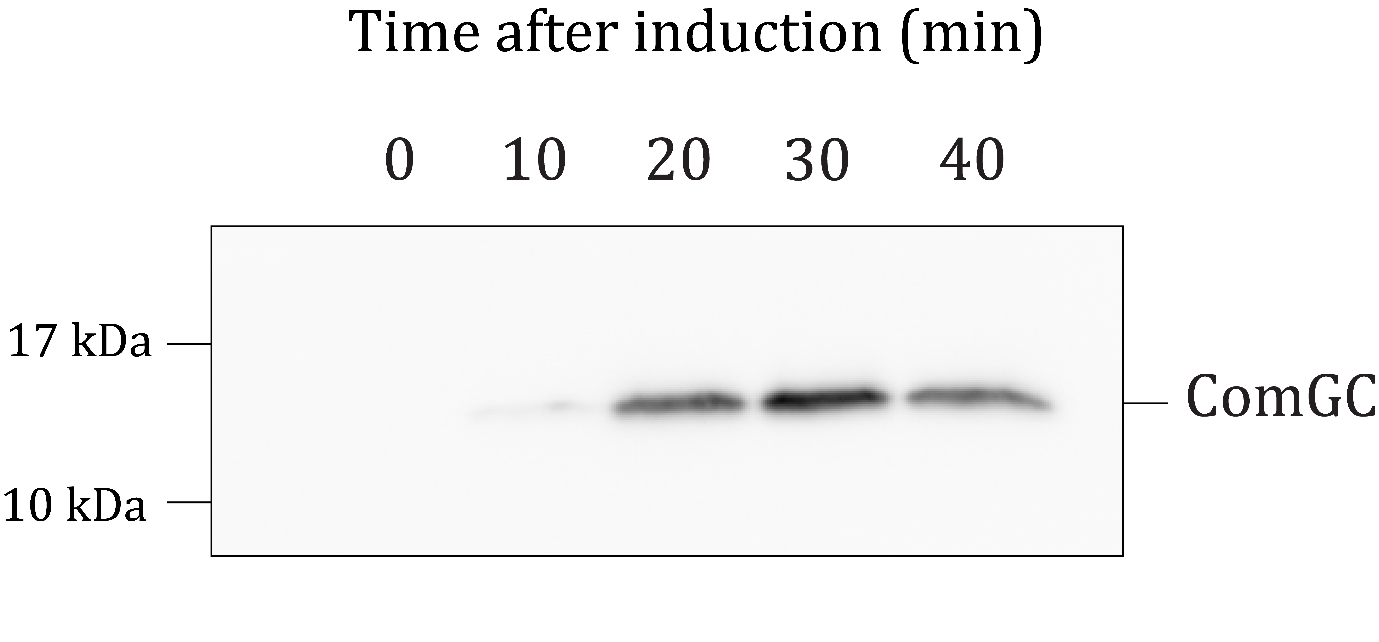

**Fig. S4** Time-series immunoblot of ComGC-FLAG to assess peak expression. Competence was induced in a 10 mL culture at OD_550_ = 0.2 and incubated for 10 to 40 minutes with increments of 10. A non-induced culture was included as a negative control (time = 0). ComGC-FLAG in samples of lysed cell extracts were detected using anti-FLAG antibodies. The strongest signal was detected 30 minutes after competence induction.

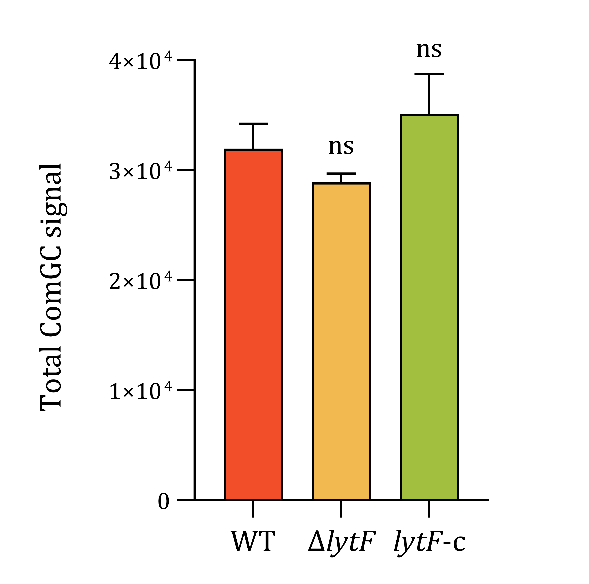

**Fig. S5** Effect of LytF on total levels of ComGC-FLAG of the wild-type (RM32), a *lytF*-knockout (RM33), and a *lytF*-complementation strain (RM125). The ComGC-FLAG signal intensities from immunoblots of extracellular and intracellular fractions were measured using ImageJ (Schneider et al., 2012) and summarised to estimate the total relative ComGC-FLAG amount. The difference in means between the strains was determined to be statistically non-significant using a One-way ANOVA and Dunnett’s test (p > 0.05).
